# Automatic Body Morphometric Analysis of Adult Zebrafish Using MicroCT

**DOI:** 10.1101/2025.09.19.677450

**Authors:** Mohammed Kanani, Ronald Young Kwon

## Abstract

Zebrafish (*Danio rerio*) are an important model for the study of musculoskeletal genetics, development, disease, regeneration, and evolution. While micro-computed tomography (microCT) allows for the assessment of quantitative 3D morphometric measures related to body size and shape and lean mass in adult zebrafish, computing these measures is typically a labor-intensive process. The purpose of this study was to develop fully automatic methods for the calculation of lean volume, fineness ratio, anterior and posterior swim bladder length, and standard length from microCT scans of adult zebrafish. Here, we show that built-in functions in the open-source software FIJI/ImageJ can be combined into fully automated methods to compute these measures. We show that measures computed by automated methods compare favorably to those computed using manual and semi-automated methods while significantly improving the throughput of data collection and eliminating inter-observer variability. We have implemented these methods as ImageJ macros, providing an accessible tool to facilitate the use of microCT for body morphometric analyses in adult zebrafish.

## INTRODUCTION

Zebrafish (Danio rerio) are an important model for the study of musculoskeletal genetics, development, disease, regeneration, and evolution. Their favorable experimental attributes, such as small size, high fecundity, low cost, and genetic tractability, make zebrafish particularly well-suited for rapid throughput studies. Micro-computed tomography (microCT) has emerged as a valuable modality for high-resolution, non-invasive analysis of the adult zebrafish musculoskeletal system, enabling the detailed study of complex morphology of bone and lean tissue in three dimensions [1-7]. In order to fully realize the potential of microCT-based phenotypic screens in zebrafish, there is a need for semi- and fully automatic image analysis pipelines.

Our lab and others have demonstrated the potential to quantify morphometric measures related to body size and shape and lean mass in microCT scans of adult zebrafish [8, 9]. Lean volume, which is defined as the volume of all voxels not classified as air, bone, or fat in microCT scans, has been shown to be closely related to skeletal muscle mass and thus serves as an indicator of skeletal muscle accrual [3, 8]. The fineness ratio, measured as the ratio of standard length to dorsoventral height, is a measure of body shape that correlates with swimming speed in some fishes [10]. The swim bladder is an air-filled organ that develops from the foregut endoderm and contributes to buoyancy regulation; the lengths of the anterior and posterior swim bladder chambers therefore provide an indicator of its development and functional capacity, which is important when assessing genetic models of swim bladder disorders or toxicological studies [11]. Finally, standard length, which is defined as the length from the tip of the snout to the base of the caudal fin rays, is an important indicator of growth and developmental progress [12].

Computing morphological measures from microCT scans is usually a labor- and time-intensive process. In our prior studies, fineness ratio, swim bladder length, and standard length were evaluated using manually-placed landmarks. The placement of these landmarks is not only tedious but is also prone to inter-observer variability. For the computation of lean volume, our lab previously described a semi-automated procedure [8]. However, one limitation of this method was the need for users to manually inspect thresholds for each fish and make adjustments to the threshold to appropriately segment the desired tissue. Moreover, users were required to interface with multiple pieces of software and feed in outputs from one piece of software into the other, which was not conducive to rapid throughput screens where hundreds or even thousands of scans are processed.

The purpose of this study was to develop fully automatic methods for the calculation of lean volume, fineness ratio, anterior and posterior swim bladder length, and standard length from microCT scans of adult zebrafish. Here, we show that built-in functions in the open-source software FIJI/ImageJ [13] can be combined into fully automated methods to compute these measures. We show that these automated methods produce values that compare favorably to manual and semi-automated methods while significantly improving throughput and avoiding inter-observer variability. We have implemented these methods as ImageJ macros, providing an accessible tool to facilitate the use of microCT for body morphometric analyses in adult zebrafish.

## MATERIALS AND METHODS

### Dataset

The dataset for this study consisted of 131 whole-body microCT scans of adult wildtype and somatic mutant (crispant) zebrafish. For full details on the generation and scanning of animals, see methods related to animal processing and scanning described in Watson et al [8]. Briefly, we analyzed crispants and their wildtype clutchmates for five genes: *wnt16* (22 crispant; 24 control fish), *fam3c* (14 crispant; 14 control fish), *tspan12* (14 crispant; 15 control fish), and *ing3* (14 crispant; 14 control fish). In our studies, similar results were observed for wildtype and crispant animals for all genes, and thus we did not separate animals by genotype in our final analyses.

All studies were performed on an approved protocol in accordance with the University of Washington Institutional Animal Care and Use Committee (IACUC# 4306–01). Studies were conducted in mixed sex animals. All fish were generated from the same genetic background (wildtype AB strain). Fish were sacrificed at 90 days postfertilization. Fresh or frozen fish were wrapped in a strip of paper towel and scanned using a vivaCT40 MicroCT scanner (Scanco Medical, Switzerland), with 21 µm isotropic voxel resolution, 55kVp, 145 mA, 1024 samples, 500proj/180°, 200 ms integration time [4]. DCM image stacks (folders containing DCM images, each corresponding to a slice in the stack) were generated using the manufacturer’s software.

Each DCM image stack contained a single fish with slices along the stack running in the anterior-posterior direction, with the first slice located at the tip of the snout, and the last slice ending approximately at the posterior end of the caudal fin bone rays. Moreover, in each DCM image stack, background voxels (i.e., voxels not containing tissue) were pre-segmented and set to zero. Note that DCMs must have a similar format for methods described below to function appropriately.

### Automated Calculation of Lean Volume

An ImageJ macro was developed to automate the calculation of lean volume (Figure 1). The script prompts the user to select the directory containing the microCT scans or a group of related scans. Once the directory is selected, a corresponding CSV file is created. A DCM series of an adult fish is loaded into ImageJ, followed by thresholding with two global histogram techniques, ‘Default’ and ‘MaxEntropy’. We have previously shown that the ‘Default’ threshold effectively provides a lower threshold to distinguish air from lean tissue, while ‘MaxEntropy’ provides an upper threshold to distinguish lean from bone tissue [8]. Threshold calculations are performed on the slice corresponding to 40% along the stack (referred to henceforth as the 1/2.5th slice), as we have previously shown this to consistently capture a slice containing bone, lean tissue, and the swim bladder, facilitating threshold calculations [8]. Thresholds are applied across the entire stack. For each slice, the area of lean tissue is computed and cumulatively summed. This sum is then multiplied by the voxel size to calculate the total lean volume. Key parameters, including the number of slices in the scan, the slice where thresholds were computed, the lower and upper thresholds computed using the ‘Default’ and ‘MaxEntropy’ algorithms, and the total lean volume, are recorded in the CSV file. This process is repeated for all scans in the directory.

**Figure 1:**
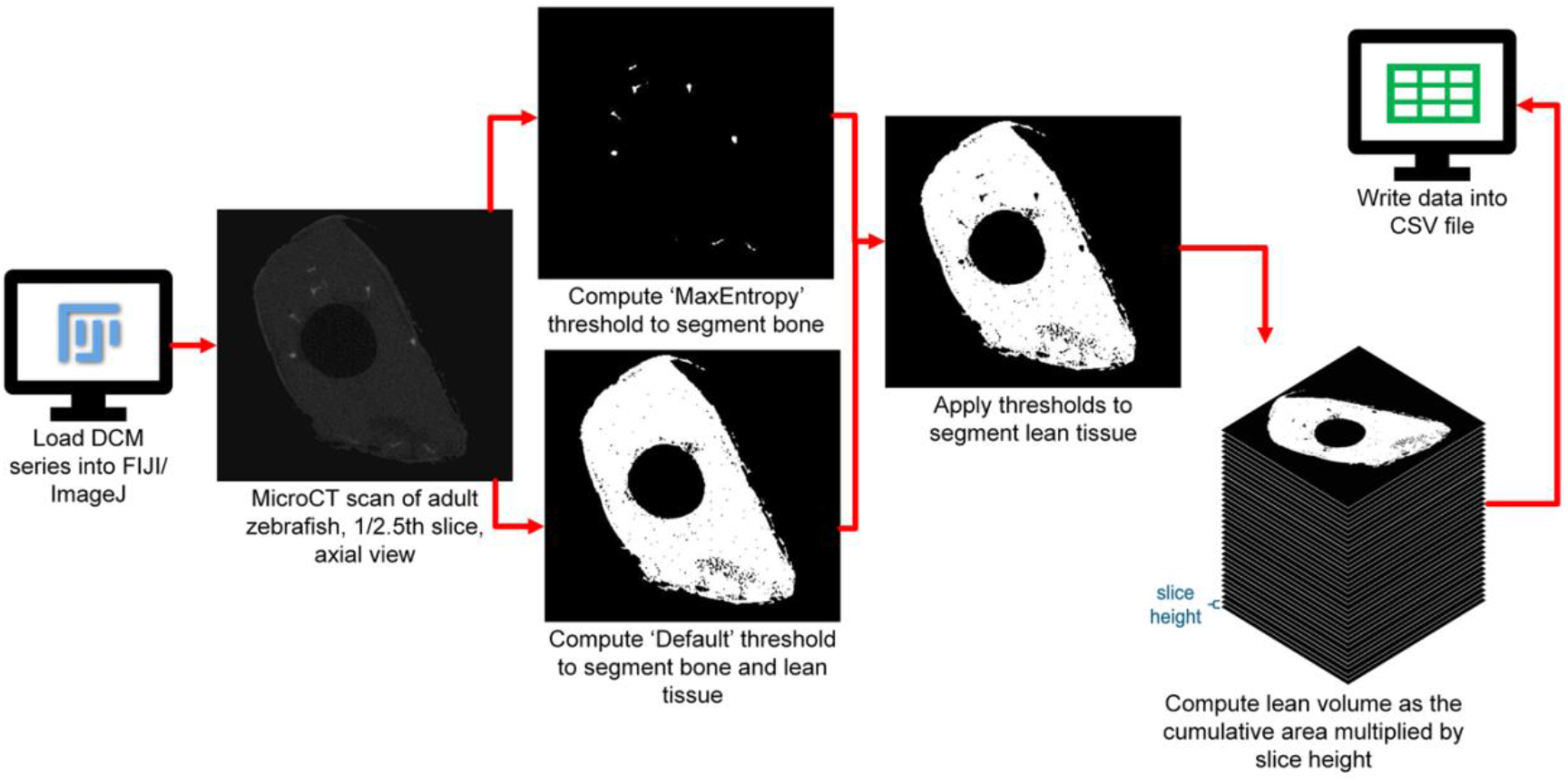
Procedure for calculation of lean volume. In this procedure, thresholds are automatically computed and used to segment lean tissue, and the total lean volume is determined by summing the areas of lean tissue across slices and multiplying by the slice height.

### Automated Calculation of Standard Length, Dorsoventral Height, and Fineness Ratio

#### Standard Length

To compute standard length, we developed and tested two different algorithms (Figure 2). In both methods, our approach was to identify the slice in the stack containing the base of the caudal fin rays. Note that in our images, the first slice was located at the tip of the jaw, and thus we did not need to develop an algorithm to identify this landmark.

**Figure 2:**
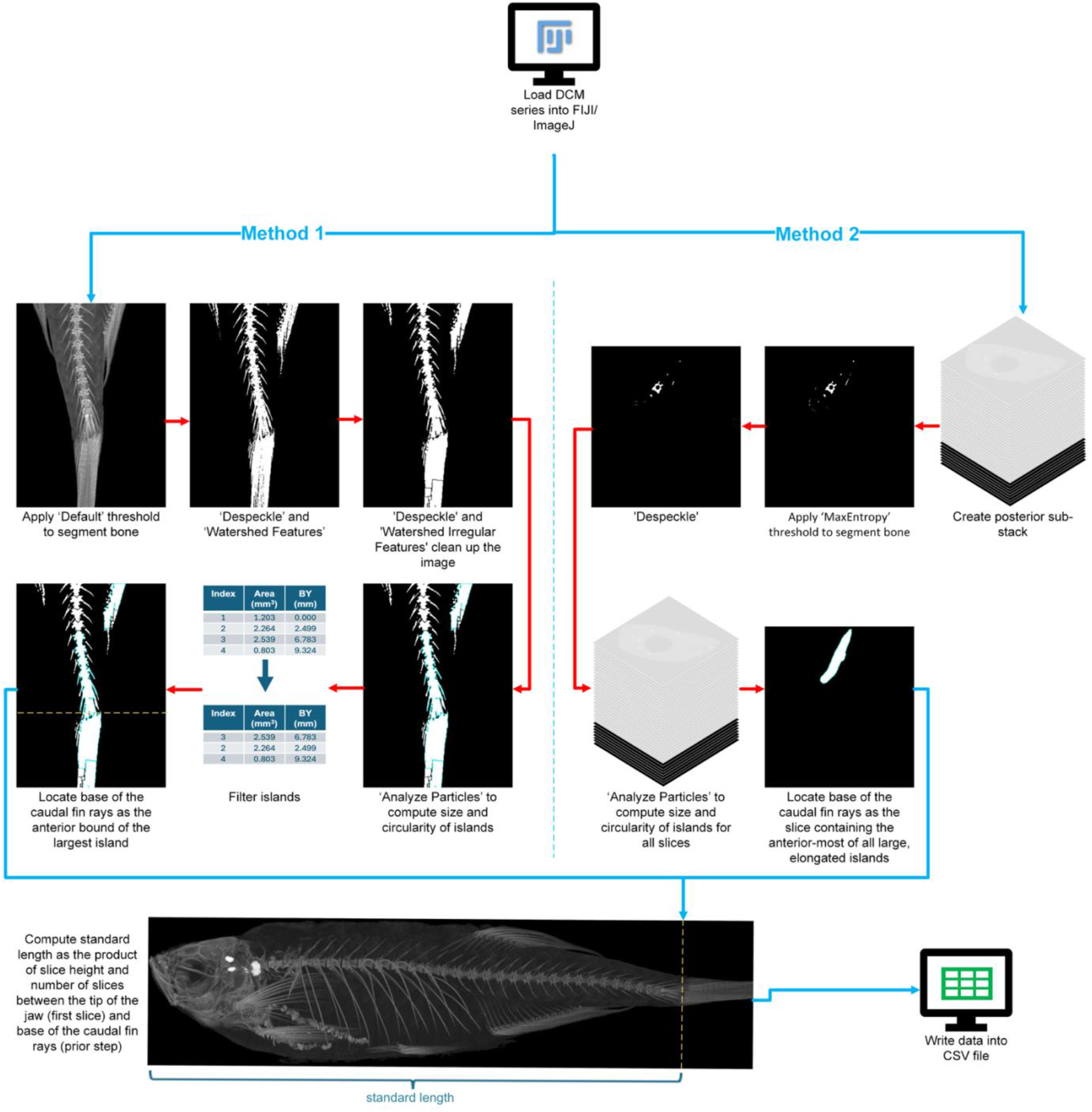
Procedure for calculation of standard length. In Method 1, the base of the caudal fin rays is located by analyzing sagittal maximum intensity projections. Here, after bone segmentation, the ‘Watershed Irregular Features’ tool is used to separate the caudal vertebrae from the caudal fin rays, the latter of which appear as one large island due to partial volume effects. The location of the base of the caudal fin rays is determined by the anterior-most boundary (BY) of the largest island in the region. In Method 2, the base of the caudal fin rays is located by traversing through the stack. Specifically, after bone segmentation, the ‘Analyze Particles’ tool is used to determine the size and elongation of islands in each slice; the location of the base of the caudal fin rays is determined as the slice containing the anterior-most of structures meeting a certain size and elongation.

Method 1 locates this region in the maximum intensity projection and takes advantage of the fact that the caudal fin rays appear as a single large “island” (i.e., connected component) of segmented pixels after thresholding due to partial volume effects. First, a sub-stack of the posterior two-thirds of the image stack is created to narrow the region of interest (ROI) where the base of the caudal fin rays can be identified. A sagittal maximum intensity (‘MaxIntensity’) projection of this sub-region is then thresholded using the ‘Default’ method to segment the bone voxels. The ‘Despeckle’ tool is applied to remove noise, followed by the ‘Watershed Irregular Features’ tool to separate the hypurals from the caudal fin rays. Empirically, this method was found to perform more reliably than the default ‘Watershed’ tool. Next, the ‘Analyze Particles’ function is used to assess the morphology of these islands, and filtered based on size and circularity. Specifically, the algorithm searched for particles at least 0.40 mm^2^ and a circularity of 0.05-1.00 (defined as 4π(area/perimeter^2^) such that a value of 1.00 resembles a perfect circle). In this way, islands with rod- or needle-like contours are excluded. Particles in the top fifth of the stack that interface with the ROI border are likely to correspond to the dorsal or anal fins and were also excluded. The areas and bounds of the identified islands are recorded and sorted by size. The largest island’s anterior bound is taken as the start of the caudal fin, and the slice containing it is recorded.

Method 2 locates the base of the caudal fin rays by again taking advantage of the fact that the caudal fin rays appear as a single large island of segmented pixels after thresholding. However, rather than doing so in a maximum intensity projection as in Method 1, it instead locates this region by surveying the stack along the anterior-posterior axis. After loading the volume, a sub-stack of the posterior-most quarter of the stack is created. Once the sub-stack is created, the first slice in the sub-stack is analyzed. A ‘MaxEntropy’ threshold is applied to segment bone pixels. The images are then processed with the ‘Despeckle’ tool, followed by ‘Analyze Particles’ to identify the anterior-most island exceeding 0.35 mm^2^ with a circularity of 0.05-0.50, corresponding to the start of the caudal fin rays. This approach is based on the observation that this region consistently presents as a larger, elongated structure, distinct from the smaller, more irregular vertebral shapes located more anteriorly. After identifying the caudal fin ray base, the slice index is recorded. For both methods, once the slice containing the base of the caudal fin rays is identified, this number is multiplied by the voxel size to compute standard length.

#### Dorsoventral Height

To compute dorsoventral height, we generated a sagittal maximum intensity projection, applied a ‘Huang’ threshold to segment the body, and applied the ‘LocalThickness’ tool (Figure 3). The dorsoventral height was computed as the maximum thickness value within the region.

**Figure 3:**
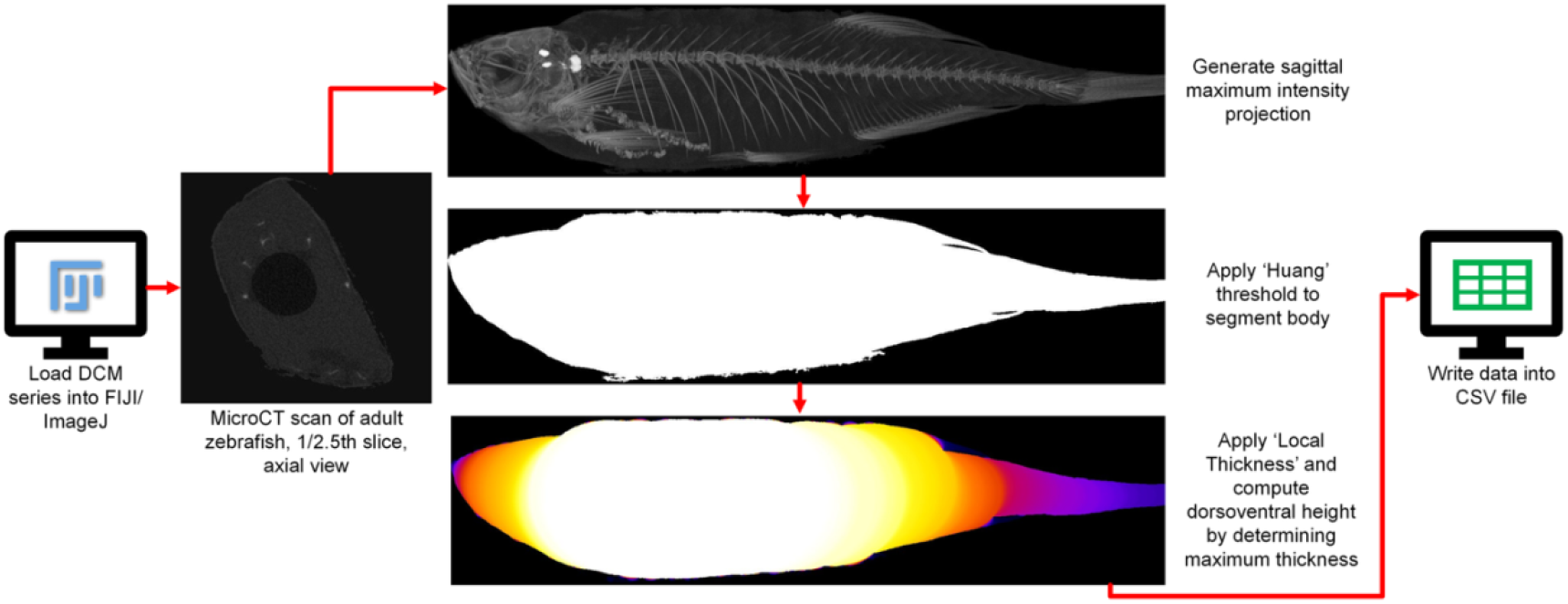
Procedure for calculation of dorsoventral height. In this procedure, the fish body is segmented in a sagittal maximum intensity projection, the ‘Local Thickness’ tool is applied, and the dorsoventral height is determined as the maximum thickness.

#### Fineness Ratio and Macro

Once automated methods for computing standard length and dorsoventral height were developed, we streamlined their calculation by creating scripts. The fineness ratio was computed as the ratio of standard length to dorsoventral height. All methods were implemented in ImageJ macros.

### Automated Calculation of Swim Bladder Chamber Lengths

Finally, we developed an algorithm to compute the length of the anterior and posterior swim bladder chambers and implemented this within an ImageJ macro (Figure 4). After prompting the user to select a working directory, the process begins by generating an axial average intensity (‘AvgIntensity’) projection to determine the fish’s orientation relative to the horizontal axis. The fish is then rotated to align the swim bladder with the horizontal axis. A minimum intensity (‘MinIntensity’) projection is applied to isolate the swim bladder. At this stage, the anterior and posterior bladder compartments are visible but often appear to be joined, and the image is noisy,so the ‘Despeckle’, ‘Remove Outliers’, and ‘Watershed Irregular Features’ tools are used to separate them. The ‘Analyze Particles’ tool is then employed to identify the bladder compartments and measure their lengths based on the major axis Feret diameter of each particle.

**Figure 4:**
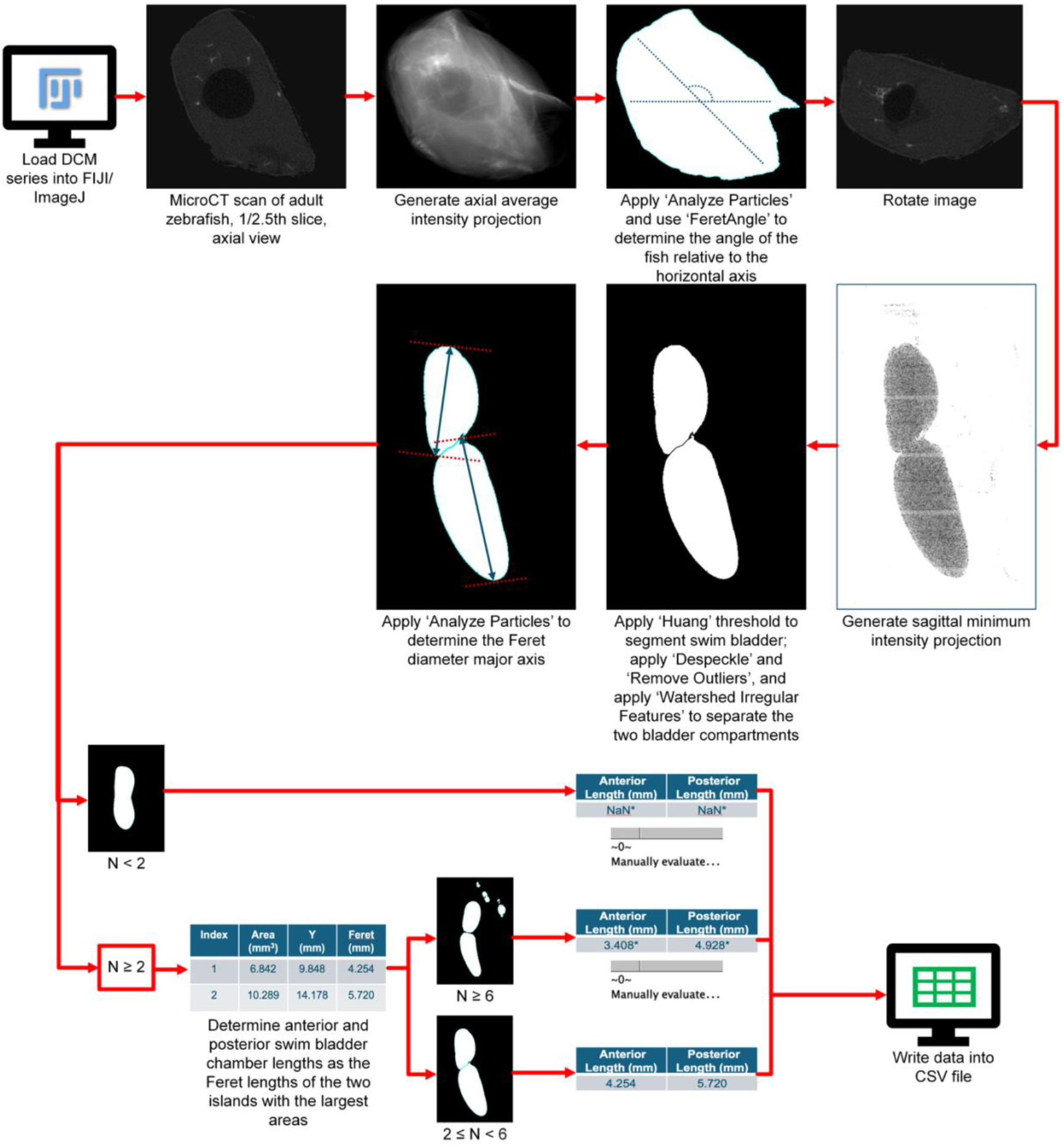
Procedure for calculation of swim bladder lengths. Here, images are rotated, the swim bladders are segmented in a sagittal minimum intensity projection, and the anterior and posterior swim bladder lengths are computed as the major axes of the two largest islands. Results are output in a CSV file, with manual evaluation suggested for uncertain cases.

If fewer than two particles are detected, computation is aborted, and the script records both bladder lengths as ‘NaN’ in the CSV file. A message is displayed to the user, advising a manual evaluation of the scan. If at least two particles are found, the two largest areas are assigned as the anterior and posterior bladder compartments based on their positions. However, when the particle count exceeds six, distinguishing the swim bladder compartments becomes challenging in our experience. In such cases, a message is generated suggesting manual evaluation, but the measurements are still recorded with an asterisk to indicate uncertainty. For particle counts between two and six, the measurements are recorded without annotation. The anterior and posterior swim bladder lengths are recorded into a CSV file. This process is repeated for all scans in the directory.

## Analysis

To evaluate method performance, we assessed the R^2^ correlation between values computed using automated (test method) and manual or semi-automated (comparative method) methods. We also generated Bland-Altman plots to qualitatively evaluate the agreement between the two measurement methods. All statistical analyses were performed in GraphPad Prism 10.

## Data Availability

The ImageJ macros are included as Supplemental Data. We have also included a folder containing sample microCT scans used in this study to demonstrate their operation.

## RESULTS AND DISCUSSION

### Lean Volume

We first compared lean volume in 131 fish quantified using automated methods to semi-automated methods we previously described. The semi-automated methods described previously were nearly identical to the automated methods developed here, with the exception that users (i) interfaced with multiple pieces of software by feeding in outputs from one piece of software into the other, and (ii) manually changed the slice used for threshold calculation if the 1/2.5^th^ slice did not contain a sufficient amount of lean, bone, and swim bladder. As expected, comparison of lean volume quantified using automated and semi-automated methods showed a high correlation (R^2^ = 0.9993) (Figure 5A). The mean difference in values (i.e., bias, automated – semi-automated) was -0.98 mm^3^, with 95%CI of the difference in values (i.e., limits of agreement, LOA) being -1.3 to -0.68 mm^3^. Expressed as a percentage of averaged values, this equated to a small positive mean bias of -0.8% with LOA of -4.3% to 2.6%. Inspection of the Bland-Altman plot suggested a lower scatter in the differences as the magnitude of lean volume increased, as well as a tendency for values of the difference to decrease with increasing magnitude (Figure 5B).

**Figure 5:**
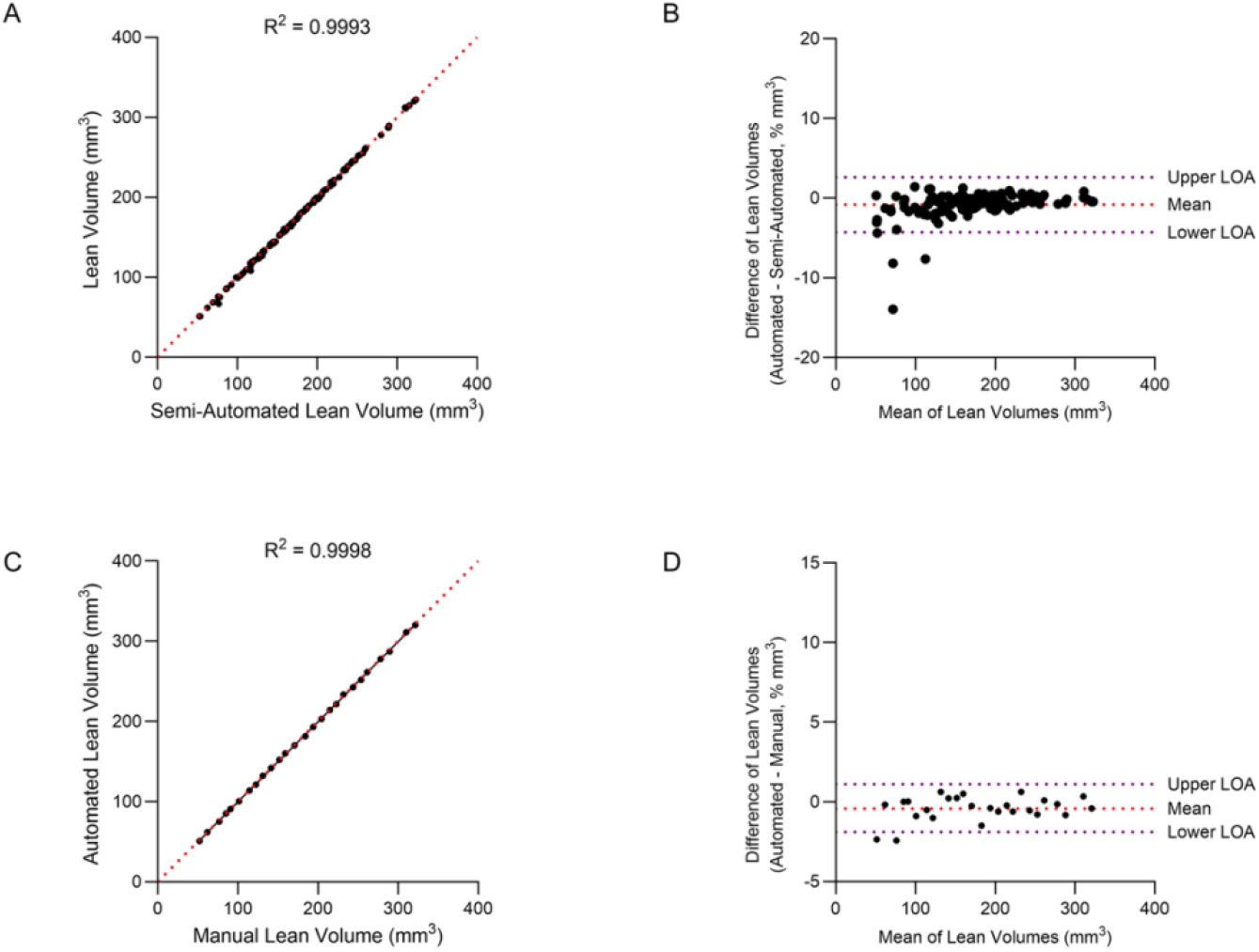
Comparison of lean volume computed using automated and semi-automated or manual methods. Scatterplots (left column) and Bland–Altman plots (right column) compare values computed using automated and semi-automated (1^st^ row) or manual (2^nd^ row) methods.

Next, we compared the automated method for measuring lean volume to values obtained manually. For this, we selected a subset of 26 fish and used manually selected thresholds to segment lean tissue. When comparing lean volume quantified using automated and manual methods, correlation remained high (R^2^ = 0.9998) (Figure 5C). The mean difference in values (automated – manual) was -0.58 mm^3^ (−0.4%), with LOA calculated to be -1.0 to -0.15 mm^3^ (−1.9% to 1.1%). There was no obvious change in the scatter in the differences or in the values of the difference with increasing magnitude in the Bland-Altman plot (Figure 5D). These results suggest that the automated method for quantifying lean volume demonstrates good agreement with both semi-automated and manual methods.

### Standard Length, Dorsoventral Height, and Fineness Ratio

We developed and tested two methods for quantifying standard length. Using Method 1, values quantified using automated methods and manual methods showed a high correlation (R^2^ = 0.9579) (Figure 6A). The mean difference in values (automated – manual) was -0.18 mm (−0.9%) with LOA calculated to be -0.27 to -0.084 mm (−5.6% to 3.8%). When quantifying standard length using Method 2, values quantified using automated methods and manual methods showed a high correlation (R^2^ = 0.9708) (Figure 6C). The mean difference in values (automated – manual) was -0.05 mm (−0.3%) with LOA calculated to be -0.13 to 0.029 mm (−4.4% to 3.8%). For both Method 1 and 2, Bland-Altman plots revealed no obvious change in the scatter in the differences or in values of the difference with increasing magnitude (Figure 6B, 6D).

**Figure 6:**
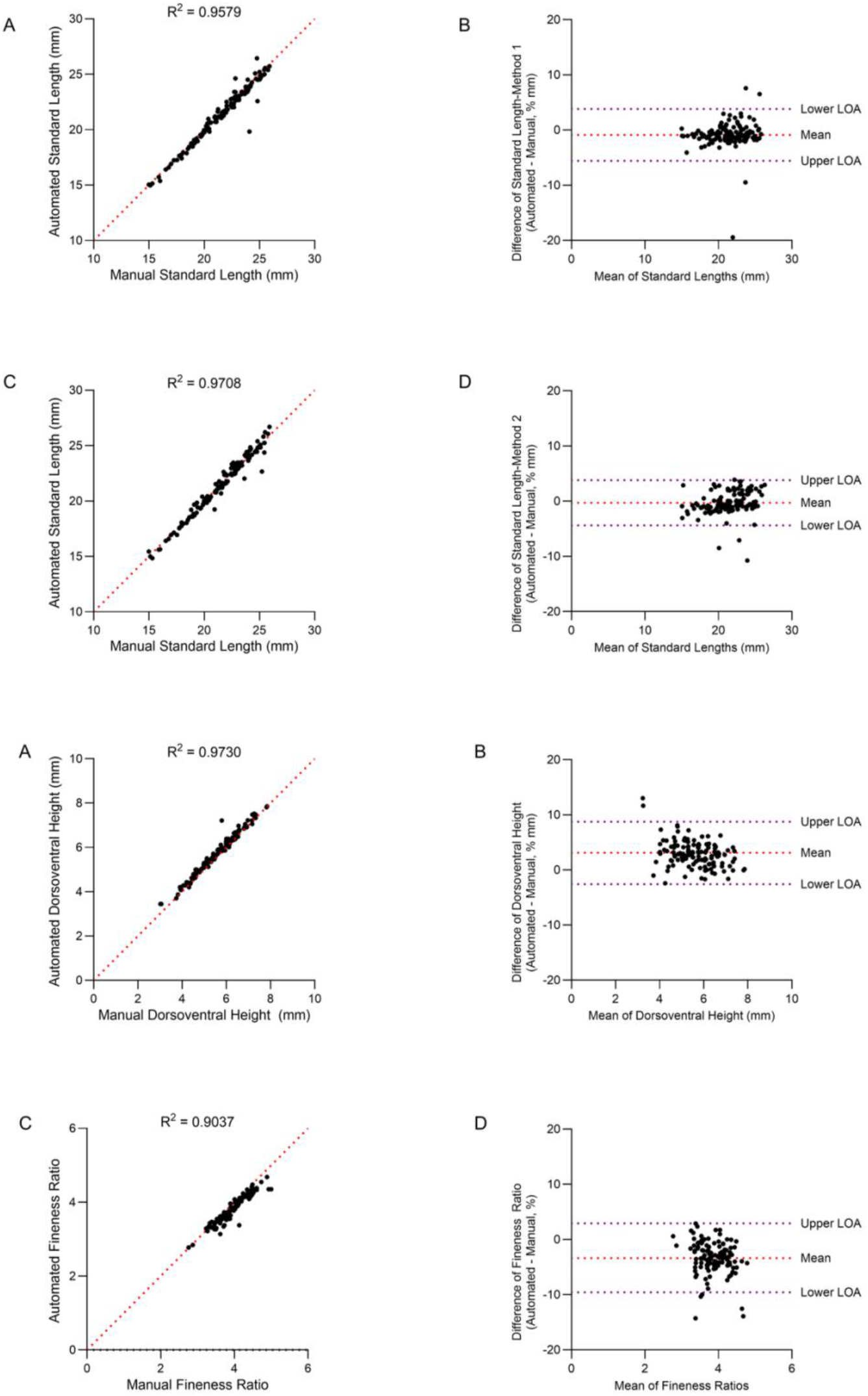
Comparison of standard length, dorsoventral height, and fineness ratio computed using automated and manual methods. Scatterplots (left column) and Bland– Altman plots (right column) compare values computed using automated and manual methods. Shown are results for standard length using Method 1 (1^st^ row), standard length using Method 2 (2^nd^ row), dorsoventral height (3^rd^ row), and fineness ratio (4^th^ row).

We compared the automated method for measuring dorsoventral height to values obtained manually. Comparison of dorsoventral height using automated and manual methods showed a high correlation (R^2^= 0.9730) (Figure 6E). The mean difference in values (automated – manual) was 0.17 mm (3.1%), with LOA calculated to be 0.14 to 0.19 mm (−2.6% to 8.7%). There was no obvious change in the scatter in the differences or in values of the difference with increasing magnitude in the Bland-Altman plot (Figure 6F).

For fineness ratio (standard length/dorsoventral height), comparison of values obtained using automated and manual methods showed a high correlation (R^2^ = 0.9037) (Figure 6G), albeit lower than for standard length or dorsoventral height alone. The mean difference in values (automated – manual) was -0.13 (−3.4%), with LOA calculated to be -0.15 to -0.11 (−9.6% to 2.9%). Inspection of the Bland-Altman plot revealed no obvious change in the scatter in the differences or in values of the difference with increasing magnitude (Figure 6H).

### Swim Bladder Lengths

When evaluating swim bladder lengths, of the 131 scans analyzed, 121 (92%) returned a confident measurement. For these 121 scans, comparison of anterior swim bladder length using automated and manual methods showed a high correlation (R^2^ = 0.9898) (Figure 7A). The mean difference in values (automated – manual) was 0.04 mm (1.1%), with LOA calculated to be 0.02 to 0.05 mm (−1.7% to 4.0%). For posterior swim bladder length, values quantified using automated methods and manual methods showed a high correlation (R^2^ = 0.9817) (Figure 7C). The mean difference in values (automated – manual) was -0.08 mm (−2.3%) with LOA calculated to be -0.09 to -0.06 mm (−7.4% to 2.7%). For both anterior and posterior swim bladder lengths, there was no obvious change in the scatter in the differences with increasing magnitude in the Bland-Altman plot; however, there was a tendency for values of the difference to decrease with increasing magnitude (Figure 7B, 7D).

**Figure 7:**
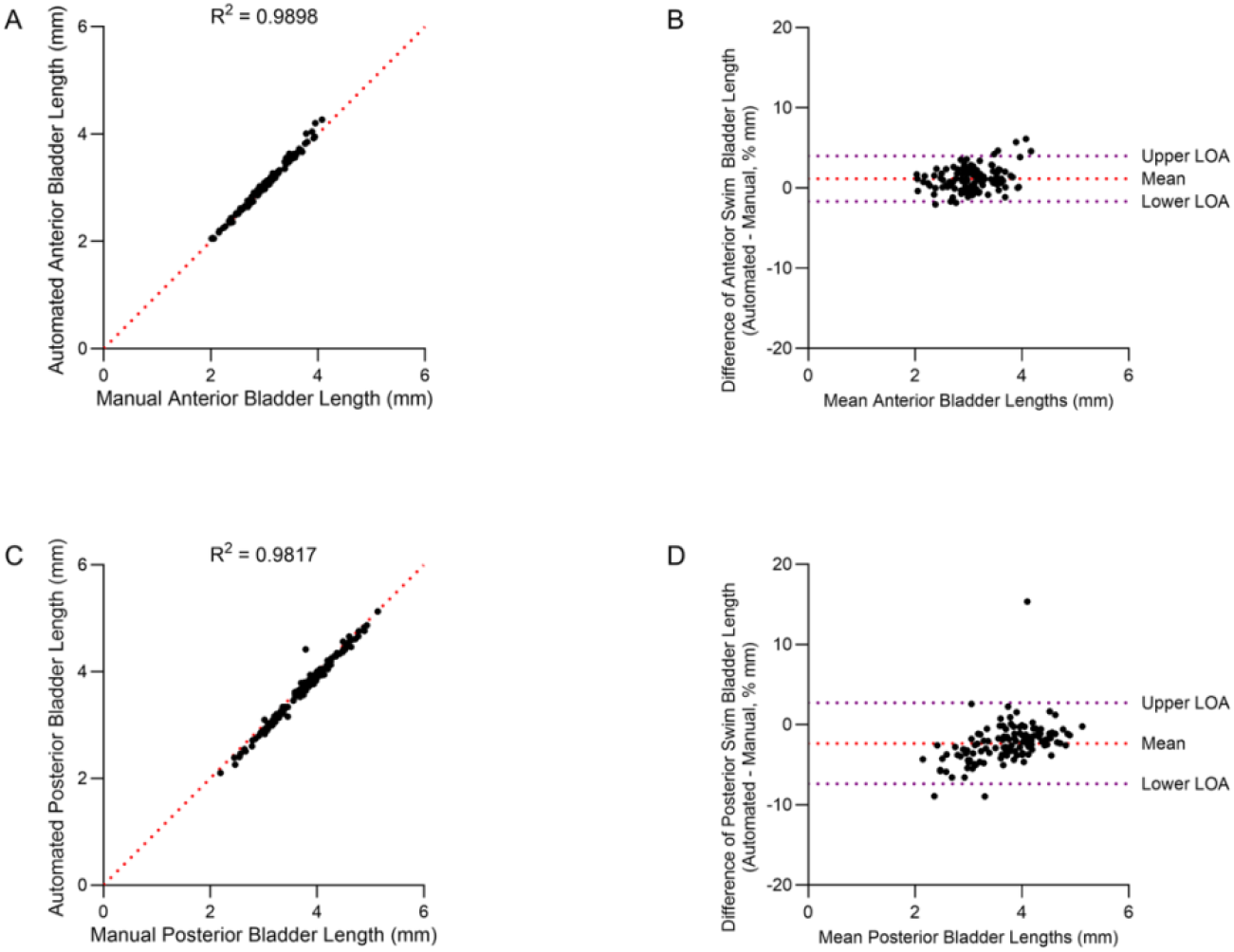
Comparison of swim bladder chamber lengths computed using automated and manual methods. Scatterplots (left column) and Bland–Altman plots (right column) compare values computed using automated and manual methods. Shown are results for anterior swim bladder chamber length (1^st^ row) and posterior swim bladder chamber length (2^nd^ row).

### Application to Different Datasets Should Be Carefully Evaluated

Although the automated methods performed well in the datasets tested here, their application to different datasets should be carefully evaluated. For instance, when calculating lean volume, thresholds are calculated on the 1/2.5th slice, as we empirically found that this slice contains lean tissue, bone, and swim bladder to facilitate the calculation of thresholds. However, during testing, we found that the ‘Default’ threshold did not segment lean tissue and bone effectively if the swim bladder is not present in this slice. While the good agreement between automated and semi-automated/manual methods suggests that this was relatively rare, this could pose challenges for fish of different ages or stages of development. Similarly, for swim bladder chamber lengths, in approximately 10% of our test set, anterior and posterior swim bladder compartments could not be confidently identified. Again, variations in animal size, anatomy, and mounting could influence the ability to distinguish anterior and posterior swim bladder compartments and thus should be tested on a case-by-case basis. Finally, differences in experimental conditions could also influence methods for the measurement of standard length. Both Method 1 and 2 identify the base of the caudal fin rays by taking advantage of the fact that the caudal fin rays appear as a single large island of segmented pixels after thresholding due to partial volume effects. However, this feature may not be conserved when analyzing high-resolution scans where partial volume effects are minimal. It is noteworthy that both methods produced similar agreement with manual methods; while Method 2 had slightly higher agreement compared to Method 1, this method also requires longer computation time. Thus, when choosing the method to be used for an application, throughput and agreement with manual methods should be considered on a case-by-case basis.

## Conclusions

In summary, we have developed a set of automated methods for assessing measures of adult zebrafish body morphology using microCT. Overall, we observed good agreement between values obtained using automated and semi-automated or manual methods, with R^2^ correlations ranging from 0.9037 to 0.9998, and values of mean bias ranging from -3.1% to 3.4%. The expected benefits of these methods include improved speed and efficiency, as well as increased reproducibility due to the removal of inter-observer variability. Although the automated methods performed well in the datasets tested here, their application to different datasets should be carefully evaluated. In particular, the use of fish other genotypes, ages/developmental stages, mounting methods, and microCT scanning instruments/acquisition settings could influence method performance. We have implemented these methods as macros in the open-source software ImageJ to facilitate broad accessibility, as well as to facilitate modifications in different experimental settings. Thus, this study provides a potentially useful tool for researchers using zebrafish for musculoskeletal research, such as in the context of rapid genetic screens or other applications requiring the analysis of large numbers of microCT scans.

## ACKNOWLEDGMENTS

Research reported in this publication was supported by the National Institute of Arthritis and Musculoskeletal and Skin Diseases of the National Institutes of Health under Award Number AR074417 and the Ernest M. Burgess Endowed Chair for Orthopaedic Investigation.

