## Supplementary material for "Automatic Body Morphometric Analysis of Adult Zebrafish Using MicroCT": measureLeanVolume.ijm

```
// Define the root folder
rootDir = getDirectory("Choose the root directory");

// Get the list of folders in the root directory
list = getFileList(rootDir);

// Create or open the CSV file
csvFile = rootDir + "leanVolume.csv";
File.open(csvFile);
File.append("DCM Name,Total Slices,(1/2.5)th Slice,Default Threshold,MaxEntropy Threshold, Volume (mm3)", csvFile);

// Run script for all DCMs in directory
// Process DCM file or DCM folder
for (i = 0; i < list.length; i++) {
	// DCM File
    if (isDICOMFile(rootDir + list[i])) {
        // Process each folder
        processDicomFiles(rootDir + list[i]);
    }
    // DCM Folder
    if (File.isDirectory(rootDir + list[i])) {
        // Process each folder
        processDicomFiles(rootDir + list[i]);
    }
}

print("Completed!");

// Computes Lean Volume
function processDicomFiles(folder) {
    // Open the DICOM series
    //run("Image Sequence...", "open=[" + folder + "] sort");
	open(folder);

    // Get the number of slices
    totalSlices = nSlices();

    // Calculate the specific slice
    sliceIndex = round(totalSlices / 2.5);

    // Go to the specific slice
    setSlice(sliceIndex);

    // Apply default threshold and get the top value
    setAutoThreshold("Default dark");
    getThreshold(lower, upper);
    defaultThreshold = lower;

    // Apply MaxEntropy threshold and get the top value
    setAutoThreshold("MaxEntropy dark no-reset");
    getThreshold(lower, upper);
    maxEntropyThreshold = lower;
    
    setThreshold(defaultThreshold, maxEntropyThreshold);
    run("Convert to Mask", "background=Dark black");

    run("Clear Results");   // First, clear the results table
    run("Set Measurements...", "area mean integrated shape limit");
  
    // loop through each slice in the stack. Start at n=1 (the first slice), 
    // keep going while n <= nSlices (nSlices is the total number of slices in the stack)
    // and increment n by one after each loop (n++)
    for (n=1; n<=nSlices; n++) {  
       setSlice(n);  // set the stack's current slice to n
       run("Measure");   // Run the "Measure" function in ImageJ
    }

    // Create a variable to store the area measured in each slice
    totalArea = 0;
    
    // Loop through each result from 0 (the first result on the table) to nResult (the total number of results on the table)
    for (n=0; n < nResults; n++)
    {
       totalArea += getResult("Area",n);   // Add the area of the current result to the total
    }
    
    // Get the calibration information from ImageJ and store into width, height, depth, and unit variables. 
    getVoxelSize(width, height, depth, unit);
    // Calculate the volume by multiplying the sum of area of each slice by the depth
    volume = totalArea*depth;
    

	// Extract the folder name from the full path
    folderName = getFolderName(folder);

	// Append data to the CSV file
    File.append(folderName + "," + totalSlices + "," + sliceIndex + "," + defaultThreshold + "," + maxEntropyThreshold + "," + volume, csvFile);

    // Close the image
    close();
}

// Function to extract the folder name from the full path
function getFolderName(path) {
    parts = split(path, "/");
    return parts[parts.length - 1];
}

// Function to check if a file is a .dcm file
function isDICOMFile(file) {
    return endsWith(file, ".DCM");  // Checks if the file name ends with .dcm
}
```
