## Supplementary material for "Automatic Body Morphometric Analysis of Adult Zebrafish Using MicroCT": measureLengthHeightV1.ijm

```
// Define the root folder
rootDir = getDirectory("Choose the root directory");

// Get the list of folders in the root directory
list = getFileList(rootDir);

// Create or open the CSV file
csvFile = rootDir + "fishLengthHeight.csv";
File.open(csvFile);
File.append("DCM Name, Length (mm), Height (mm), Fineness Ratio", csvFile);


// Run script for all DCMs in directory
// Process DCM file or DCM folder
for (i = 0; i < list.length; i++) {
	// DCM File
    if (isDICOMFile(rootDir + list[i])) {
        // Process each folder
        processDicomFiles(rootDir + list[i]);
    }
    // DCM Folder
    if (File.isDirectory(rootDir + list[i])) {
        // Process each folder
        processDicomFiles(rootDir + list[i]);
    }
}

print("Completed!");

// Computes Length/Height
function processDicomFiles(DCM) {
	open(DCM);
	fileName = getTitle();
	
	// Returns DCM dimensions
	getVoxelSize(width, height, depth, unit);
	totalSlices = nSlices();
	twoThirds = round(2/3*totalSlices);
	stackLength = depth*totalSlices;

	// Computes height using LocalThickness, saves as fishHeight
	run("Rotate 90 Degrees Left");
	run("Reslice [/]...", "output=0.021 start=Top avoid");
	run("Z Project...", "projection=[Max Intensity]");
	setAutoThreshold("Huang dark");
	run("Convert to Mask");
	
	run("Local Thickness (masked, calibrated, silent)");
	getStatistics(area, mean, min, max, std, histogram);
	
	close();
	close();
	close();
	
	fishHeight = max;

	// Computes fish length
	
	// Create substack
	run("Make Substack...", "slices=" + twoThirds + "-" + totalSlices);
	run("Reslice [/]...", "output=0.021 start=Top avoid");
	run("Z Project...", "projection=[Max Intensity]");
	setAutoThreshold("Default dark");
	run("Convert to Mask");
	
	subLength = getHeight()* depth;

	// Process stack
	run("Despeckle");
	run("Watershed Irregular Features", "erosion=1 convexity_threshold=0.83 separator_size=0-Infinity");
	
	run("Set Measurements...", "area slice bounding redirect=None");
	run("Clear Results");   // First, clear the results table
	run("Analyze Particles...", "size=0.40-Infinity circularity=0.05-1.00 display clear overlay");
	
	nParticles = nResults;  // Number of particles
	
	areas = newArray(nParticles);
	yPositions = newArray(nParticles);

	// Extract measurements (major, minor axis, and Y-position)
	for (i = 0; i < nParticles; i++) {
	    areas[i] = getResult("Area", i);    // Area
	    yPositions[i] = getResult("BY", i);  // Y-coordinate (position)
	}

	// Initialize largest value and its index
	largestValue = 0;
	largestIndex = 0;

	// Find the largest value and its index
	for (i = 0; i < areas.length; i++) {
	    if (areas[i] > largestValue && yPositions[i] > 0.2*subLength) {
	        largestValue = areas[i];  // Update largest value
	        largestIndex = i;  // Update index of largest
	    } 
	}
	
	yTop = yPositions[largestIndex];
	fishLength = stackLength - (subLength - yTop);
	
	
	// Computes fineness ratio
	fineness = fishLength/fishHeight;

	// Append data to the CSV file
	File.append(fileName + "," + fishLength + "," + fishHeight + "," + fineness, csvFile);

	close("*");
}

// Function to extract the folder name from the full path
function getFolderName(path) {
    parts = split(path, "/");
    return parts[parts.length - 1];
    
    
// Function to check if a file is a .dcm file
function isDICOMFile(file) {
    return endsWith(file, ".DCM");  // Checks if the file name ends with .dcm
}
```
