## Supplementary material for "Automatic Body Morphometric Analysis of Adult Zebrafish Using MicroCT": measureSwimBladder.ijm

```
// Define the root folder
rootDir = getDirectory("Choose the root directory");

// Get the list of folders in the root directory
list = getFileList(rootDir);

// Create or open the CSV file
csvFile = rootDir + "swimBladderMetrics.csv";
File.open(csvFile);
File.append("DCM Name, Anterior Length (mm), Posterior Length (mm)", csvFile);

// Run script for all DCMs in directory
// Process DCM file or DCM folder
for (i = 0; i < list.length; i++) {
	// DCM File
    if (isDICOMFile(rootDir + list[i])) {
        // Process each folder
        processDicomFiles(rootDir + list[i]);
    }
    // DCM Folder
    if (File.isDirectory(rootDir + list[i])) {
        // Process each folder
        processDicomFiles(rootDir + list[i]);
    }
}

print("Completed!");

// Computes Swim Bladder Metrics
function processDicomFiles(DCM) {

	open(DCM);
	fileName = getTitle();

	// Process DCM
	run("Rotate 90 Degrees Left");
	run("Reslice [/]...", "output=0.021 start=Top avoid");
	run("Z Project...", "projection=[Min Intensity]");
	setAutoThreshold("Huang");
	setOption("BlackBackground", true);
	run("Convert to Mask");
	run("Despeckle");
	run("Remove Outliers...", "radius=20 threshold=50 which=Bright");
	close("\\Others");
	run("Watershed Irregular Features", "erosion=1 convexity_threshold=0.97 separator_size=0-Infinity");
	
	run("Set Measurements...", "area centroid bounding fit feret's stack limit redirect=None decimal=3");
	run("Clear Results");   // First, clear the results table
	run("Analyze Particles...", "size=50-Infinity pixel display exclude clear overlay");


	// Get the particle measurements
	nParticles = nResults;  // Number of particles
	
	if (nResults < 2){
		print("Manually evaluate " + fileName);
		File.append(fileName + "," + "NaN" + "," + "NaN", csvFile);
	} else {
	
		// Initialize arrays to store the measurements
		majors = newArray(nParticles);
		minors = newArray(nParticles);
		areas = newArray(nParticles);
		yPositions = newArray(nParticles);
		
		
		// Extract measurements (major, minor axis, and Y-position)
		for (i = 0; i < nParticles; i++) {
		    majors[i] = getResult("Feret", i);  // Major axis
		    minors[i] = getResult("MinFeret", i);  // Minor axis
		    areas[i] = getResult("Area", i);    // Area
		    yPositions[i] = getResult("Y", i);  // Y-coordinate (position)
		}
		
		// Initialize variables for the largest and second largest values
		largestValue = areas[0];
		secondLargestValue = -1; // Start with a value smaller than any in the array
		
		largestIndex = 0;
		secondLargestIndex = -1;
		
		// Find the largest value and its index
		for (i = 1; i < areas.length; i++) {
		    if (areas[i] > largestValue) {
		        secondLargestValue = largestValue;  // Update second largest
		        secondLargestIndex = largestIndex;  // Update index of second largest
		        largestValue = areas[i];  // Update largest value
		        largestIndex = i;  // Update index of largest
		    } else if (areas[i] > secondLargestValue) {
		        secondLargestValue = areas[i];  // Update second largest value
		        secondLargestIndex = i;  // Update index of second largest
		    }
		}
		
		
		// Get the Y-position of the two largest particles
		yTop = yPositions[largestIndex];
		ySecond = yPositions[secondLargestIndex];
		
		
		// Determine anterior and posterior labels based on Y-position
		if (yTop > ySecond) {
		    anterior = majors[secondLargestIndex];  // Uppermost particle
		    posterior = majors[largestIndex];  // Lowermost particle
		} else {
		    anterior = majors[largestIndex];  // Uppermost particle
		    posterior = majors[secondLargestIndex];  // Lowermost particle
		}
		
		
		if (nResults >= 6){
			print("Manually evaluate " + fileName);
			File.append(fileName + "," + anterior + "*" + "," + posterior + "*", csvFile);
		} else {
			// Append data to the CSV file
		    File.append(fileName + "," + anterior + "," + posterior, csvFile);
		}
	
	}

	close("*");

}


// Function to extract the folder name from the full path
function getFolderName(path) {
    parts = split(path, "/");
    return parts[parts.length - 1];
    
    
// Function to check if a file is a .dcm file
function isDICOMFile(file) {
    return endsWith(file, ".DCM");  // Checks if the file name ends with .dcm
}
```
