## Supplementary material for "Automatic Body Morphometric Analysis of Adult Zebrafish Using MicroCT": README.md

# Automatic Body Morphometric Analysis of Adult Zebrafish Using MicroCT — ImageJ/Fiji Macros

This repository provides four Fiji/ImageJ macros for body morphometric analysis using microCT scans of adult zebrafish, as well as sample DICOM images. Each macro loads DICOM data (either a single `.dcm` stack file or a folder of DICOM slices), processes the volume, and exports measurements to CSV. Full algorithms and validation are described in the accompanying paper.

## Macros

- `measureLeanVolume.ijm` — computes lean-tissue volume  
  CSV columns: `Total Slices`, `(1/2.5th Slice)`, `Default Threshold`, `MaxEntropy Threshold`, `Volume (mm3)`
- `measureLengthHeightV1.ijm` & `measureLengthHeightV2.ijm` — compute standard length, dorsoventral height, and fineness ratio  
  CSV columns: `Standard Length`, `Height`, `Fineness Ratio`
- `measureSwimBladder.ijm` — measures anterior and posterior swim bladder lengths  
  CSV columns: `Anterior Length`, `Posterior Length`

> The macros accept either a single `.dcm` stack or a folder containing DICOM slices.

### Example data (DICOM)
The `DCM/` folder includes four example stacks:
- `example_fish_1.dcm`, `example_fish_2.dcm` — single-file DICOM stacks  
- `example_fish_3/`, `example_fish_4/` — folders of individual DICOM slices  

We have included example data as both single-file DiCOM stacks as well as folders of individual DICOM slices in order to demonstrate the ability of the macros to accept either format. In this data set, we have intentionally included a DICOM stack (example_fish_2.dcm) where the anterior and posterior swim bladder chambers are not readily identified. When run with measureSwimBladder.ijm, it will skip measurement and return an error code to prompt manual evaluation. This demonstrates how the macro handles cases when the required conditions for analysis are not met.


---

## Requirements

- Fiji / ImageJ (tested with ImageJ 2.16.0 / 1.54p; Java 8)
- BioVoxxel Toolbox (installed via Fiji Update Sites)
- IJBlob library (`ij-blob.jar`) on Fiji’s classpath

---

## Install BioVoxxel Toolbox (macOS & Windows)

1. Open Fiji → Help → Update…  
2. Click Advanced mode → Manage update sites  
3. Check BioVoxxel → Close → Apply changes → Restart Fiji  
   - Docs: https://imagej.net/plugins/biovoxxel-toolbox  
   - How to follow an update site: https://imagej.net/update-sites/following

> After restart, you should see BioVoxxel under Plugins.

---

## Install IJBlob (`ij-blob.jar`)

Download the JAR (choose one source):
- ImageJ plugin page: https://imagej.net/plugins/ij-blob  
- Releases / mirrors: https://sourceforge.net/projects/ijblob/files/  
- Direct JAR listing (example): https://sites.imagej.net/Biomedgroup/jars/ij_blob-1.4.9.jar-20160714123114

### macOS
1. Quit Fiji.  
2. Copy the downloaded `ij-blob.jar` to:  
   `/Applications/Fiji.app/jars/`  (preferred)  
   (Alternatively: `/Applications/Fiji.app/plugins/`)  
3. If Gatekeeper flags the file, clear quarantine in Terminal:  
   ```bash
   xattr -d com.apple.quarantine "/Applications/Fiji.app/jars/ij-blob.jar"
   ```  
4. Restart Fiji.  
5. Verify: Plugins → Utilities → List JARs → confirm `ij-blob*.jar` is listed.

### Windows
1. Close Fiji.  
2. Copy `ij-blob.jar` to:  
   `C:\Fiji.app\jars\`  (preferred)  
   (Alternatively: `C:\Fiji.app\plugins\`)  
3. If Windows blocked the download: right-click the JAR → Properties → check Unblock → OK.  
   (PowerShell alternative:)  
   ```powershell
   Unblock-File -Path "C:\Fiji.app\jars\ij-blob.jar"
   ```
4. Restart Fiji.  
5. Verify: Plugins → Utilities → List JARs → confirm `ij-blob*.jar` is listed.

> If you previously saw `java.lang.NoClassDefFoundError: ij/blob/ManyBlobs`, installing IJBlob resolves it.

---

## Quick Start (all macros)

1. Fiji → Plugins → Macros → Run… → select one of the `.ijm` files.  
2. When prompted, select either:
   - a single `.dcm` stack (e.g., `DCM/example_fish_1.dcm`), or
   - a folder of DICOM slices (e.g., `DCM/example_fish_3/`).  
3. Choose an output location when prompted. A CSV is saved with the measurements.  
4. Open the CSV in your spreadsheet/editor.

Notes
- Physical units come from DICOM metadata (PixelSpacing, SliceThickness). If missing, set the scale first: Analyze → Set Scale…  
- For large stacks, increase Fiji memory: Edit → Options → Memory & Threads…  
- `measureLeanVolume.ijm` logs Default and MaxEntropy Auto Threshold values for transparency.

---

## Troubleshooting

- BioVoxxel menu missing → Re-run Help → Update… → Manage update sites, ensure BioVoxxel is checked → Apply changes → restart.  
- `NoClassDefFoundError: ij/blob/ManyBlobs` → Ensure `ij-blob.jar` is in `.../Fiji.app/jars/`, then restart.  
- Empty/zero CSV → Confirm DICOM has scale metadata or set scale manually; ensure the fish is visible (window/level) and not fully thresholded out.

---
